# *Toxoplasma gondii* associates with human Benign Prostatic Hyperplasia and induces prostatic hyperplasia and accompanying urinary dysfunction in mice

**DOI:** 10.64898/2026.04.23.720409

**Authors:** Emily F. Stanczak, Tara D. Fuller, Doug W. Strand, Hanyu Xia, Oliver R. Strobel, Irene Heredero Bermejo, Gustavo Arrizabalaga, Travis J. Jerde

## Abstract

**Objectives:** Benign Prostatic Hyperplasia (BPH) is the non-cancerous enlargement of the prostate accompanied by lower urinary tract symptoms, affecting 50% of men by the age of 50^1,2^. Advanced highly symptomatic BPH exhibits large epithelial glandular nodules with microglandular/atypical adenomatous hyperplasia, but how these features form is unknown^3^. Our lab has reported that the common parasite *Toxoplasma gondii* can infect the prostate and induce glandular nodule formation in mice^3^. The objective of this study is to determine if *T. gondii* exposure in humans correlates to BPH and nodule formation and if it induces urinary dysfunction concurrent in the mouse model.

**Methods:** We assessed *Toxoplasma* exposure by serum ELISA in patients with BPH and non-BPH donor controls, and compared seropositivity rates between the groups. We further assessed the histopathology of these patients for the presence of inflammation and epithelial glandular nodule formation and compared *Toxoplasma* positive and negative samples. We determined voiding function in *Toxoplasma*-infected mice between 14 and 60 days of infection with void spot with Void Whizzard software.

**Results:** Men diagnosed with BPH are more likely to be seropositive for *Toxoplasma* than age-matched undiagnosed donor controls. In addition, BPH patients that are seropositive for *Toxoplasma* are more likely to exhibit glandular nodule formation with microglandular / adenomous hyperplasia than seronegative *BPH patients*. In animal studies, *Toxoplasma* infection results in abnormal void patterns concurrent with microglandular hyperplasia and nodule formation.

**Conclusions:** These results suggest that *Toxoplasma* may be contributing to BPH pathology and lower urinary tract dysfunction in both humans and mice, opening new insights into the development of this important disease. The results also serve to further characterize this model of prostatic hyperplasia and define it as a potential urinary dysfunction model.

## INTRODUCTION

Benign Prostatic Hyperplasia (BPH) is the non-cancerous enlargement of the prostate, often accompanied by lower urinary tract symptoms (LUTS)^1^. Approximately 50% of men by the age of 50 will develop BPH symptoms, and serious complications, including frequency, urgency, and pain can arise from long-term BPH-LUTS^1^. Currently available therapeutics treat the symptoms of BPH; however, these therapies are non-curative and are associated with side-effects that can be therapy-limiting^1^. Additionally, patients can develop therapeutic resistance or respond differently to available therapies drugs resulting in a treatment gap that leaves many men in the advanced stages of BPH with few to no treatment options available^1,2^.

BPH typically presents as widespread inflammation throughout the transition zone of the prostate^3^. Early-stage BPH prostates exhibit stromal nodules of non-glandular tissue and developing fibrosis and epithelial reactive hyperplasia^3^. In contrast, advanced specimens commonly exhibit growing epithelial, glandular nodules in the transition zone. These nodules consist of epithelial hyperplasia of small glands with adenomous features termed atypical adenomatous hyperplasia, adenosis, or microglandular hyperplasia^3^. These epithelia consist of well-circumscribed, crowded, small to medium-sized glands with discontinuous but clearly present basal cell layer. This layer joins with neighboring small epithelia to form defined nodular regions of epithelium inside a defined fibrotic stromal compartment^3^. Nodules make up 2-20% of advanced BPH tissue and are distinguished from prostate cancer by their clear cytoplasm, distinct basal cell layer, fusion with benign glands, presence of corpora amylacea, and lack of oncogene expression^3^.

Several factors have been proposed to contribute to BPH including genetics, metabolic disease, diet, shift in testosterone: estrogen ratio, and bacterial, viral, or parasitic infections^1,2^. Interestingly, the common household parasite *Toxoplasma gondii* can infect mouse prostates and cause similar pathology^4^. *Toxoplasma* is an obligatory intracellular protozoan parasite that can infect any nucleated cell within practically all warm blooded animals^5,6^. Importantly, *Toxoplasma* can establish a chronic latent infection that infects approximately 30% of the world and 20% of the United States^5,6^. Although most infections are asymptomatic, severe symptoms and even death can occur in immunocompromised individuals and fetuses infected in utero. While the chronic infection has long been considered latent in immunocompetent hosts, it is now understood that a chronic inflammatory state can result from infection in several organs. Here, we show that human *Toxoplasma* infection correlates with BPH diagnosis and glandular nodule formation. We also demonstrate that mice infected with *Toxoplasma* exhibits abnormal void patterns, and develop microglandular hyperplasia and glandular nodule formation.

## METHODS

### Sample Collection

Human blood serum and prostate specimens were collected from BPH patients receiving resection or prostatectomy for BPH at UT Southwestern Medical Center and placed in the UTSW biorepository with full patient consent and in accordance with the ethical standards of the Declaration of Helsinki. Control donor specimens were collected from deceased male organ donors whose families consented at the Southwest Transplant Alliance under IRB STU 112014-033. All BPH diagnoses were made by urologists based upon prostate exam, symptom score, and outlet obstruction. Further confirmation of BPH histopathology was done by pathologists at UTSW.

### Serological Analysis of Human Blood Samples

Serology testing for *Toxoplasma* IgG antibodies was completed using the *Toxoplasma* IgG ELISA Immunoassay following the manufacturer’s instructions (BIO-RAD, Hercules, CA, USA) and measured on a Biotek Synergy H1 Hybrid Reader.

### H&E Staining and Pathology Scoring

Paraffin-embedded human prostates were sectioned to 5 microns, stained with H&E using standard protocols, and imaged with Leica scope 2500 and Aperio ImageScope 12.3.3. Analysis was completed blindly using Image J. Sections were scored based on a scoring system previously developed (Fig S1, 2)^8^.

For analysis of inflammation, three 20x magnified images from H&E-stained prostate samples were selected from random locations and evaluated. High inflammatory infiltrate (HII) is deemed any area of over 200 leukocytes. Fields scored as not present (0) have no evidence of HII present. Fields scored as mild (1) have areas of HII less than 1 mm^2^. Fields scored as moderate (2) have areas of HII ranging between 1 and 2 mm^2^. Fields scored as severe (3) have areas of HII 2 mm^2^ or larger present.

Focality was scored as (0, not present) as no evidence of inflammation in the image evaluated, (1, focal) as inflammation in only one location of the image, (2, limited) inflammation present but in less than 25% of the image, (3, intermediate) inflammation in 25%-50% of the area evaluated, (4, widespread) inflammation in more than 50% of the area evaluated. The severity and focality scores were multiplied and the average of three separate 20x fields scores are averaged for each data point to generate the overall inflammation score as reported.

### Microglandular hyperplasia score

Three 20x magnified images from H&E-stained prostate samples were selected from random locations and evaluated for microglandular hyperplasia (MH), as defined by highly proliferative, well-circumscribed, crowded, small glands with pale to clear cytoplasm, a clear basal cell layer and minimal to no obvious cellular atypia. Areas of MH exhibit small to medium-sized nuclei and often fuse with adjacent benign glands. Intensity was scored as fields scored as not present (0) exhibit no microglandular structures; those scored as mild (1) exhibit histological confirmation of microglandular hyperplasia but no fusion into a nodule; fields scored as moderate (2) exhibit one example of MH forming into a nodule in the area evaluated; and fields scored as severe (3) exhibit more than one nodule exhibiting MH or widespread non-nodular MH.

Focality was scored as follows: not present (0) if no MH was found in the specimen; focal (1) if found one location in the section; limited (2) if found in less than 25% of the area evaluated; intermediate (3) if found in 25%-50% of the area evaluated; and widespread (4) if found in more than 50% of the area evaluated. Severity and focality scores were multiplied together for each image and a single score for each 20x field. The average of three separate 20x fields scores are averaged for each data point.

### Epithelial/Glandular Nodule Count

Nodules are defined as concentrated areas of hyperplastic glands that are tightly clustered and exhibit a papillary like infoldings, all surrounded by a well-circumscribed compact stromal surrounding that condenses stroma and becomes a separate structure from the surrounding tissue. Nodules were counted of the whole image.

### Mice Experiments

#### Animal care and infection

Male and female CBA/j, C57Bk6/j, (Jackson Laboratories) or CD-1 (Charles River) mice were housed in the Laboratory Animal Resource Center (LARC) at Indiana University School of Medicine under the National Institutes of Health Guide for the Care and Use of Laboratory Animals, after all protocols were approved by the Indiana University School of Medicine Animal Care and Use Committee (IACUC). When mice were 10-12 weeks old, they were intraperitoneally (i.p.) injected with either sterile 1X phosphate-buffered saline (PBS) or 40,000 PruΔC32 *T. gondii* tachyzoites and monitored daily for signs of infection. Pain and distress were not observed typically, however animals exhibiting overt signs such as matted fur, hunched stance, and/or overgrooming were sacrificed humanely. Surviving mice were analyzed for symptom analysis at timepoints described below, and then sacrificed by isoflurane to effect and cervical dislocation at 14, 28, and 60 DPI for molecular or pathological analysis as described below.

#### Parasite culture

*Toxoplasma gondii* parasites of the PruΔhxprt ldh2GFP (PruΔC32) strain were grown on human foreskin fibroblasts (HFFs). The HFFs were cultured in Dulbecco’s modified Eagle’s medium (DMEM—Fisher Scientific) supplemented with 10% heat-inactivated fetal bovine serum (FBS—Atlanta Biologicals), 2 mM L-glutamine, 100 U/mL penicillin, and 100 µg/mL streptomycin (Fisher Scientific) and cultures were maintained at 37 °C with 5% CO2.

#### Parasite isolation and preparation

Parasites were harvested from infected HFF cultures containing a mixture of parasites still within host cells and egressed parasites. Cultures were scraped off their flasks, passed through a 27-gauge needle to release intracellular parasites, and pelleted at 800 × g for 5 minutes at 4 °C. The parasites were washed in PBS, then pelleted again (800 × g for 5 minutes at 4 °C). The pellet was again resuspended in PBS, and the parasite concentration was determined using a hemocytometer. The parasites were prepared for infection at a concentration of 40,000 parasites/100 µL in PBS. The parasite suspension was kept on ice to maintain their viability until injection. The time from intracellular parasite release to infection was kept under 45 minutes.

### Void Pattern Analysis (VPA)

All experiments for VPA calculations were conducted between 4:00 AM and 6:00 AM during mice activity times^9^. Individual mice were removed from cages and placed into empty cages lined with Whatman chromatography paper. The mice were left in these cages in the dark for two hours, the paper was removed and replaced. After an additional two hours, the paper was removed, the mice were then returned to their shared cages. After air drying for ∼8 hours, papers were placed on a UV transilluminator, and images were taken using a Canon EOS Rebel T6 camera. Images were transferred to Photoshop and urine spots were selected and darkened for easy identification. The images were converted to binary and analyzed using the Void Whizzard software developed by the U54 O’Brien Center at the University of Wisconsin: Madison. This software measured spot size in cm using pre-set bin categories of size including 0-0.25, 0.25-0.5, 0.5-1.0, 1.0-2.0, 2.0-3.0, 3.0-4.0, and 4.0+ (cm).

### Statistical Analysis

All analyses were performed in Graphpad Prism graphing and statistics software. Summative data between multiple groups and timepoints were calculated with two-way ANOVA. Binary data for human serum analysis testing for Toxopositivity was calculated with Chi Square analysis with Yates correction.

## RESULTS

### *Toxoplasma* infection correlates with BPH diagnosis

To determine if there is a correlation between *Toxoplasma* infection and BPH, blood samples from both diagnosed BPH patients and control donors were tested for the presence of anti-*Toxoplasma* IgG. BPH patients are men who have been diagnosed by medical professionals while donors are samples from deceased patients determined to be void of BPH pathology. Of the undiagnosed donors only one tested positive for *Toxoplasma* IgG whereas 15 BPH patients tested positive for *Toxoplasma* IgG (Table 1). Due to a large gap in average age between groups, we honed our analysis to patients between the ages of 45 and 70 and age-matched them. In these age-matched groups, none of the donors tested seropositive for *Toxoplasma* IgG but 10 BPH patients tested seropositive for *Toxoplasma* IgG (Table 2). This indicates that *Toxoplasma* infection statistically associates with BPH.

**Table 1.** Seropositivity for *Toxoplasma* IgG among BPH diagnosed patients and control donors. The average age for individuals in each group is shown. Statistical analysis was completed using a chi squared analysis with a Yates’ correction; p=0.0120.

| Cohort | Average Age | No. Total Samples | No. *Toxoplasma* Sero-Positive |
| --- | --- | --- | --- |
| BPH patients | 69 | 66 | 19 |
| Non-BPH Donors | 31 | 38 | 1 |

**Table 2.** Comparison of seropositivity for *Toxoplasma* IgG between BPH diagnosed patients and control donors using only age-matched samples. Statistical analysis was completed using a chi squared analysis with a yate’s correction; p=0.0415.

| Cohort | Average Age | Total Samples | No. *Toxoplasma* Sero-Positive |
| --- | --- | --- | --- |
| *BPH* | 61 | 47 | 17 |
| *Non-BPH Donor* | 56 | 26 | 0 |

### Severity of inflammation correlates with BPH diagnosis but not *Toxoplasma* infection

Given the association between *Toxoplasma* seropositivity and BPH, we investigated whether *Toxoplasma* infection correlated with BPH pathology severity. First, we assessed inflammatory infiltrate. As expected, we found that levels of inflammatory infiltrate are increased in BPH patients compared to donors (Fig. 1A). However, there was no significant difference between infected and uninfected BPH patients (6.9 ±1.1 vs. 5.3±1.0) (Fig. 1A). This indicates that *Toxoplasma* infection does not significantly alter the levels of inflammation in the prostate.

**Figure 1.**
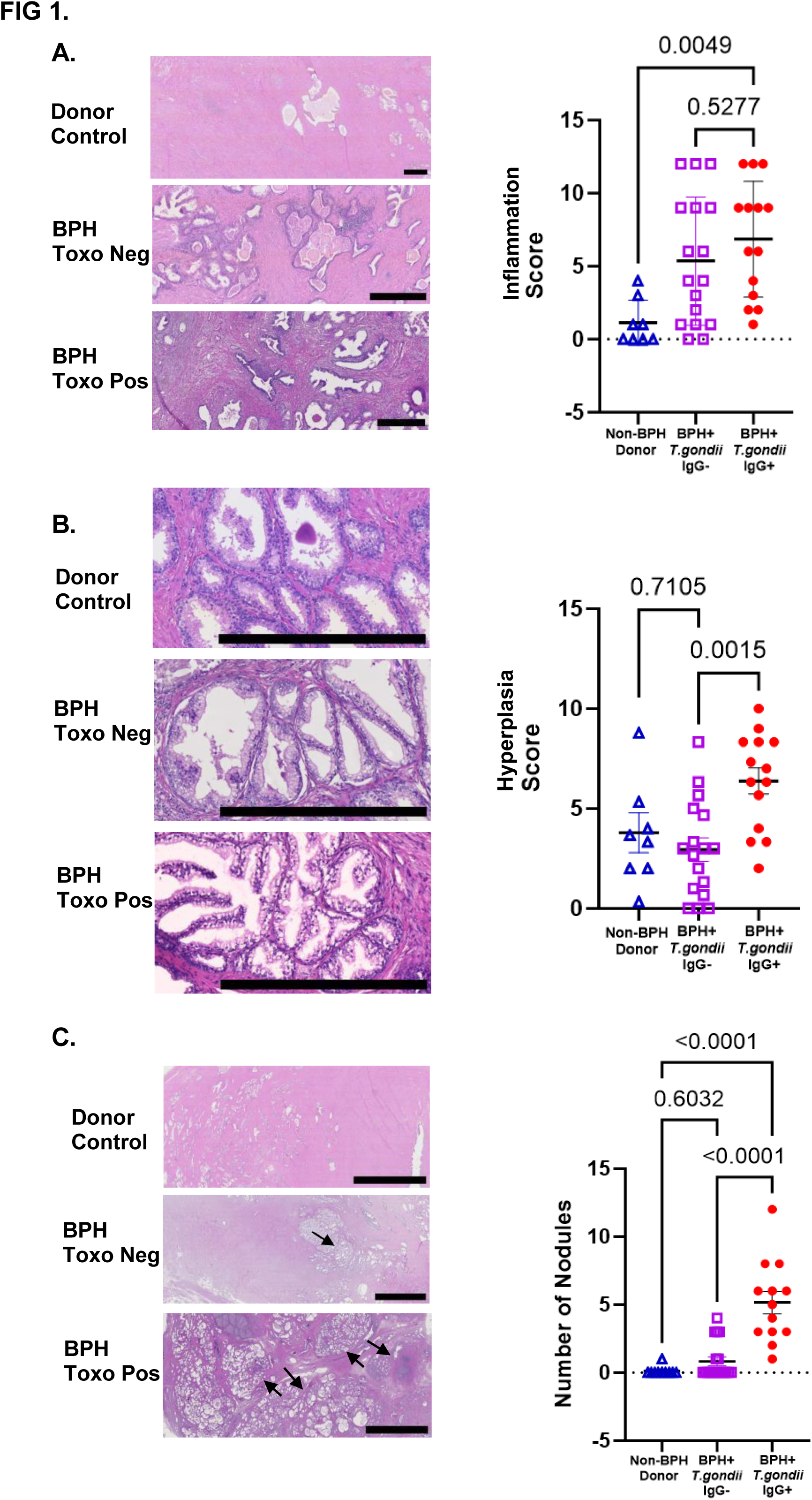
BPH histopathology in age-matched BPH diagnosed and *T*oxoplasma seropositive patients versus BPH diagnosed and *Toxoplasma* seronegative patients and donor controls. **A,** Inflammation scores were compared using a one-way ANOVA. Inflammation is more severe in both BPH and *Toxoplasma* seropositive and seronegative cohorts than in control donor samples. Graph depicts Inflammatory pathology scores in each group compared using a one-way ANOVA with images depicting each cohort. Top: Control Donor, Middle: BPH and *Toxoplasma* seronegative, Bottom: BPH and *Toxoplasma* seropositive. Scale bars indicate 500 **μ**m **B,** microglandular hyperplasia is more severe and advanced in age-matched BPH and *Toxoplasma* seropositive patient samples than BPH and *Toxoplasma* seronegative patients and control donors. Graph indicates inflammatory pathology scores in each group and images depicting microglandular hyperplasia pathology scores for each category. Top: Image of control donor, Middle: BPH seronegative patient, Bottom: BPH and *Toxoplasma* seropositive patient. Scale bar is set at 500 **μ**m. **C,** Nodule presence is increased in BPH and *Toxoplasma* seropositive patient samples than BPH and *Toxoplasma* negative patient samples and control donors. Graph indicating nodule amount for each group and images depicting nodule presence in each category. Top: Image of control donor, Middle: Uninfected BPH patient, Bottom: *Toxoplasma* infected BPH patient. Scale bars indicate 5000 **μ**m.

### Microglandular hyperplasia correlates with BPH and *Toxoplasma* infection

A histological component of BPH associated with a more severe stage of the disease is microglandular hyperplasia. Interestingly, we found that microglandular hyperplasia scores of infected BPH patients were significantly higher than uninfected BPH patients and donor controls (6.4±0.6 vs. 2.9±0.6 vs 3.8±1.0 Fig. 1b). These results suggest that *Toxoplasma* infection correlates with more higher-grade hyperplastic histopathology.

### Epithelial/glandular nodule formation correlates with BPH and *Toxoplasma* infection

We found that infected BPH patients had an increased average of nodules present (5.2±0.8) compared to uninfected BPH patients (0.8±0.3) and control donors (0.1±0.1) lacking meaningful nodular hyperplasia (Fig. 1c). Interestingly, every sample from our infected BPH group presented with at least one nodule with one sample harboring 12 nodules (Table 3). These data further indicate that *Toxoplasma* infection associates with higher intensity of hyperplastic histopathology.

**Table 3.** Epithelial/glandular nodule formation scores between age-matched BPH patients seropositive and seronegative for *Toxoplasma*.

| Cohort | No. Total Samples | No. Nodular | Percent Nodular |
| --- | --- | --- | --- |
| *Toxoplasma* Positive | 17 | 17 | 100 |
| *Toxoplasma* Negative | 30 | 9 | 32 |

**Table 4.**

| Model | Inflammation | Reactive Hyperplasia | Glandular Nodules | Dysfunction | Cytokine signaling |
| --- | --- | --- | --- | --- | --- |
| *E. coli* Models | + | + | - | - | + |
| POET | + | + | - | ? | + |
| Hormonal | + | + | + | + | - |
| Cytokine OE | +/- | + | - | - | + |
| *Toxoplasma* | + | + | + | + | + |
| Human BPH | + | + | + | + | + |

### *Toxoplasma* infection in mice correlates with abnormal void patterns

Previous data revealed that infected mice develop local infection of the prostate during the acute stage of the infection as seen in H&E analysis^4^. In this study, we confirmed infection in the prostate by localization of parasitic rosettes, and that infected mice had pathology including microglandular hyperplasia and putative nodule formation ^4^ (Fig 2). Given the increased severity in hyperplasia observed in infected BPH patients and infected mice, we hypothesized that *Toxoplasma* infection correlates with increased symptomology. Accordingly, we used the mouse model to analyze the effect of infection on urinary void behavior. Three different strains were infected intraperitonially, and void patterns were recorded at 28 days post infection (d.p.i) and 60 d.p.i. *Toxoplasma*-infected C57 mice at 28 d.p.i presented with more urine spots than uninfected controls (22.1±13.5 vs. 9.4±4.3 fig.3b). This pattern is persistent at 60 d.p.i. (27.9±15.4 vs. 14.2±4.2 Fig 3). *Toxoplasma* infected CBA/J mice at 28 d.p.i also presented with more small spots compared to controls (52.0±31.8 vs. 11.7±7.5 Fig 3c). However, this pattern did not continue at 60 d.p.i. (67.5±16.5 vs. 56.8±35.0 Fig. 3c). Finally, with CD-1 mice we see an increased amount of spotting in infected mice at 28 d.p.i compared to controls (27.4±11.0 vs. 7.0±6.1 Fig. 3d). This continues at 60 d.p.i. (52.0±31.8 vs. 11.7±7.5 Fig 3d). In both CD-1 mice and C57 mice, irregular voiding behavior continues at 60. While this is also seen in CBA/J, the control group also presented a similar pattern. Nonetheless, all three strains of mouse present irregular voiding behavior at 28 d.p.i, at which timepoint the infection is transitioning from the acute to its chronic infection stage.

**Figure 2.**
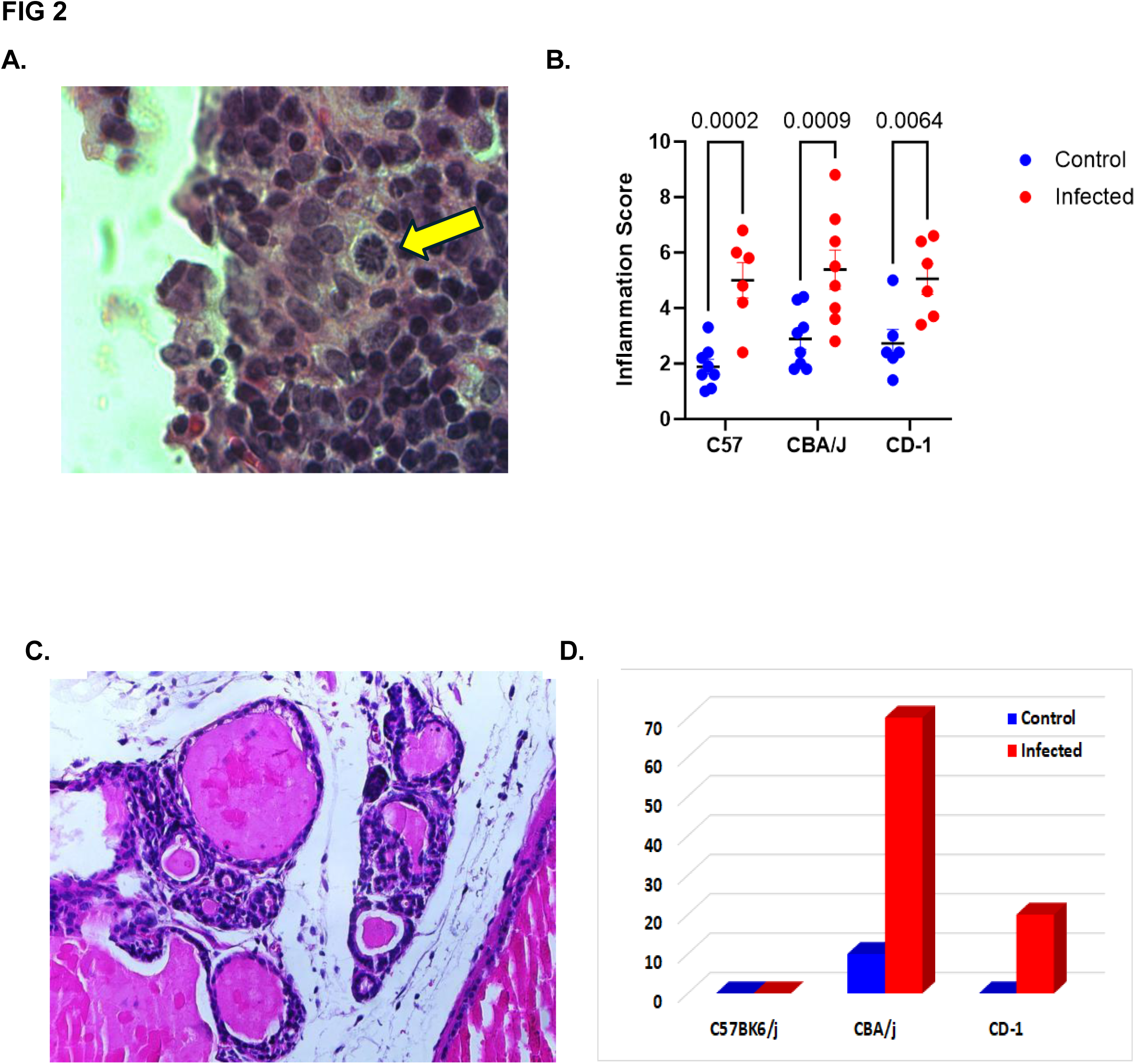
Systemic infection with *Toxoplasma* infects and inflames the mouse prostate but has mouse strain-distinct effects on hyperplasia. It has previously been shown that systemic *Toxoplasma* infects and induces inflammation in rodent prostates and induces nodular hyperplasia in the CBA/j strain of mice. The applicability of CBA/j mice to prostate modeling was unclear. To confirm this finding, we assessed infectability, inflammation, and formation of microglandular hyperplasia and nodule formation across mouse strains. **A,** a rosette of *Toxoplasma* surrounded by severe lymphocytic and monocytic inflammation in following systemic infection in CBA/j mice. **B,** Systemic *Toxoplasma* infection induces inflammation in mice in several strains (B57, CBA/J, and CD-1) as determined by H&E and crude inflammation scoring. However, only CBA/j mice exhibit a significant formation of glandular nodules **[C,D]** with microglandular hyperplasia among the strains. **C,** a region of *Toxoplasma-*associated microglandular hyperplasia and putative forming nodules; **D,** calculations of percent of mice in each strain and group that exhibit microglandular hyperplasia and/or forming nodules among mouse strains infected with *Toxoplasma* versus control. n=10, comparisons made, infected versus control in 2×2 table with Chi-square analysis with Yates correction.

**Figure 3.**
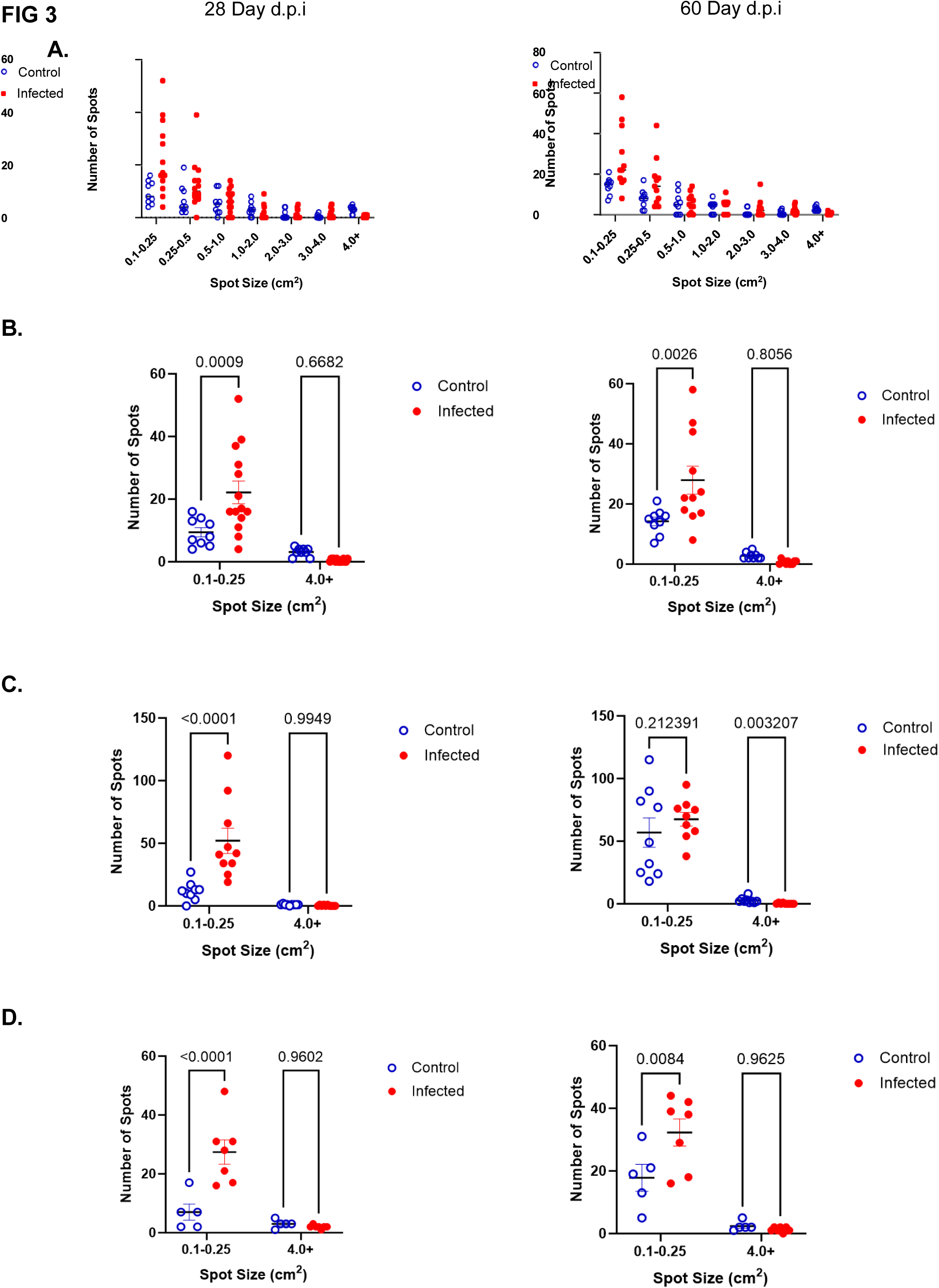
*Toxoplasma* infected mouse and uninfected control mouse number of spots counted at 28 d.p.i and 60 d.p.i in several different size categories collected. **A,** C57 mouse void pattern analysis including all spot sizes counted in the initial VoidWhizzard software analysis used **B,** C57 void pattern analysis including only the small size bin (0.1-0.25 cm) category representing spotting or irregular behavior and the larger bin size (4.0 cm) representing normal void behavior. **C,** CBA/J mouse void pattern analysis including the same spot size categories as B, and **D,** CD-1 mouse void pattern analysis including the same spot size categories as B. The left column indicates 28 d.p.i while the right indicates 60 d.p.i.

## DISCUSSION

BPH is one of the most common urological concerns in public health, yet limited information exists regarding how BPH develops^1^, and its causes and triggers. Several proposed causal factors could contribute to this multifactorial disease, but there is no single identified cause^1^. Also, it is difficult to predict BPH in individual patients, case severity, and response to treatment based upon disease history^1^. Here, we show that seropositivity to *Toxoplasma* correlates to BPH diagnosis and pathology in humans. Accordingly, our data generate the hypothesis that *Toxoplasma*, a common parasite known for causing inflammation and molecular pathway disruption in their hosts, might contribute to the development and severity of BPH in humans, as one possible trigger for inflammation that may promote prostate pathology.

Men develop symptoms and characteristics of BPH as they age regardless of *Toxoplasma* seropositivity, and the likelihood of *Toxoplasma* exposure increases with age^12,5,6^. Therefore, it is critical to consider age when comparing BPH and control individuals. We focused our seropositivity study using samples between the ages of 45-75 to ensure a similar average age in patient groups. We set 45 as the low age due to the reduced chances of BPH diagnosis prior to that age. We set 75 as the high age as most male patients develop LUTS by this age. After controlling for age, our data indicate that *Toxoplasma* exposure is associated to BPH incidence and histological hyperplasia in human prostates. Unfortunately, we were unable to control the age at which our patients were exposed to *Toxoplasma* and cannot determine if exposure time impacts pathology severity. We are also unable to account for patient lifestyle other than age, race, metabolic disorder status and BPH diagnosis status. We therefore are unable to assess exposure to cats (the definitive host for *Toxoplasma*), likelihood to have consumed contaminated produce or water or undercooked meat, geographic location, symptoms, or comorbid medical conditions. However, since *Toxoplasma* does establish a chronic infection in all hosts, it is likely that parasites are present in *Toxoplasma* positive patients at the time of tissue procurement^5,6^, and we have previously shown the parasite is present in BPH prostate specimens^4^.

A number of parasitic infections could be associated with the induction of prostatic inflammation in humans, as the immune response to protozoans in tissues is strongly and consistently T cell-driven. *Toxoplasma* was chosen for further study, primarily for two reasons: the very high prevalence of *Toxoplasma* incidence in humans across the globe, and our previous study surveying modeling systems of prostatic inflammation that found strong inflammation induction from systemic *Toxoplasma* in mice. Our study does not preclude other intracellular Th1 cell-inducing protozoan parasites being involved in inducing prostatic inflammation, including *Giardia*, *Plasmodium*, or *Cryptosporidium*. However, Toxoplasma is prevalent in up to a third of the world’s adults and infects up to 22% of Americans. The protozoan Trichomonas vaginalis has been linked to a number of urological diseases including BPH, and infects about 0.5% of the population. Given *Toxoplasma*’s prevalence, our previous study demonstrating direct prostatic inflammation induction in animal studies, and our present data in this study, *Toxoplasma* association with BPH through prostatic inflammation becomes of paramount interest as a potential inducer of prostatic inflammation.

*Toxoplasma*-positive and negative BPH patients had similarly high inflammation incidence, but widely divergent incidence of glandular nodule formations with microglandular hyperplasia. This suggests that inflammation may be a trigger for some BPH symptomology, but alone does not account for the nodule formations. There are many possible triggers for recalcitrant prostatic inflammation as men age, and systemic or local *Toxoplasma* presence is likely only one trigger. Additionally, prostate inflammation models do not typically induce glandular nodule formations, adenomous hyperplasias, and usually do not exhibit consistent LUTS.^14–17^ *Toxoplasma* in the prostate may have unique impacts on prostate pathology beyond being an inflammatory stimulus. Given that only CBA/j mice exhibit consistent nodule formation among mice, it is possible that induction of specific immune system elements specific to mouse strain may drive a different tissue response. Similarly, *Toxoplasma* may induce endocrine system effects that impact prostatic responses in a way inflammatory stimuli alone cannot^10^. Of note, only high dose hormonal modeling has been shown to induce this pathology in mice^3,10–13^. Metabolic syndrome and type II diabetes is also concurrent with BPH-LUTS and associated with prostatic inflammation, and severe prostate pathologies. Future characterization of the immune response in nodular highly symptomatic BPH-LUTS may reveal key mechanistic understandings for how specific elements of the immune response of the prostate may promote specific pathologies^18–20^. Unfortunately, we do not have symptom scores on all our patients, however all our BPH patients had sufficient symptoms to require surgical intervention, and our donor population did not have any medical record of BPH/LUTS. Therefore, we are confident in a clear difference in patient symptomology between groups.

After demonstrating the correlation between *Toxoplasma* exposure and BPH and pathological severity in humans, we turned to our mouse model to characterize further causality of *Toxoplasma* in prostate pathology^13^. Our previous work showed that inflammation and hyperplasia could be induced by systemic *Toxoplasma* infection in mice, and here we show that urinary dysfunction also is induced with infection^4^. In our study, we show that *Toxoplasma*-infected mice not only exhibit inflammation of the prostate, but this extends as a symptomatic condition in the mice, as urinary spotting is significantly increased in infected animals. In assessing the void spot analysis assay, we used spot sizes measured by Void Whizzard Image J and collected into bins of area that correspond to a range of greater than 4 cm spot to spots 0.1 to 0.25 cm^4,13^. Thus, our data showing a significant increase in spots in the 0.1-0.25 represents a significant urinary dysfunction. Additionally, we did not observe a statistically increased amount of inflammation in other organs of the urinary tract in our infected male mice, and our female mice did not show increased symptom score. Therefore, we propose that the prostate is the organ most likely to be promoting LUTS in these infected animals, though we cannot be fully certain that there is not some contribution of bladder or urethra, despite lack of obvious histopathology in these organs.

Notably, *Toxoplasma* infection induced inflammation and urinary dysfunction across all strains of mice analyzed, but only induced microglandular hyperplasia and early glandular nodule formation in the CBA/j strain. Our original study used CBA/j mice as they are common in parasite research due to their ability to harbor parasites at high titers^4^. However, as most prostate modeling occurs in C57Bk6/j and CD-1 mice, we used these strains as comparators. C57Bk6/j mice succumbed to infection titers of >1000 parasites, yet at 1000 parasites exhibited chronic prostatic inflammation. CD-1 mice sustained 40,000 parasites, showed prostatic inflammation, but failed to generate nodular formations. This indicates that voiding dysfunction as measured by number of small void spots tracks with inflammation rather than advancing glandular hyperplasia. Future studies assessing the impact of long-term hyperplasia on outlet obstruction would be warranted to determine the impact of nodular hyperplasia in the CBA/j strain, which may have a different response to the parasite. Our study highlights differences in various measures of urinary dysfunction, as well as mouse strain-specific effects of inflammation of the prostate.

## CONCLUSION

In conclusion, our data indicate that patients with BPH are more likely to have been exposed to *Toxoplasma,* and that infection with this parasite correlates with pathology in BPH patients. Additionally, mice infected with *Toxoplasma* exhibit urinary dysfunction including increased voiding of smaller volumes, which mimics what is seen in human patients with LUTS. Additionally, these mice develop microglandular hyperplasia and putative glandular nodule formations in CBA/j mice. Future studies of the immune and endocrine responses to parasitic infection may define the mechanistic understanding of these results.

## AUTHOR CONTRIBUTIONS

Drs. Emily Stanczak, Travis Jerde, Gustavo Arrizabalaga, and Doug Strand, designed experiments. Drs. Stanczak, Jerde, Fuller, and Arrizabalaga performed animal experiments with Oliver Strobel and Hanyu Xia, analyzed data and prostate pathology, and quantified staining. Dr. Strand oversaw human data collection and design. Drs. Stanczak and Jerde supervised staining quantification. Drs. Stanczak, Arrizabalaga, and Heredero Bermejo oversaw and conducted human serological testing experiments for *T. gondii*. All authors contributed to writing and reviewing the manuscript.

## COMPETING INTERESTS STATEMENT

The authors declare no relevant financial interests.

## ACKNOWLEDGEMENTS

The authors gratefully acknowledge our funding sources NIH-NIDDK 1R01DK124067 and NIH-NIAID R21AI138255-01. Additionally, the authors acknowledge the expert contributions in methodology of the U54 George M. O’Brien Center for Benign Urology Research at University of Wisconsin Madison for Void Spot Analysis and LUTS analysis in mice. The authors also greatly appreciate and acknowledge the contributions in human tissue analysis from the University of Texas Southwestern Medical School Department of Urology Biorepository for use and expert analysis of human blood and tissue specimens.

## Supplementary Information

**Supplemental table 1.** Scoring system used to quantitate inflammation and micrograndular hyperplasia in human samples.

|  |  |  | Score |
| --- | --- | --- | --- |
|  | Inflammation | Microglandular Hyperplasia |  |
| **Not Present** | No evidence of inflammatory infiltrate areas in tissue | Only pseudostratified epithelium in the area evaluated | 0 |
| **Mild** | Area of inflammatory infiltrate found less than 1 mm<sup>2</sup> | Evidence of epithelial layering in at least one location juxtaposed to inflammation in area evaluated | 1 |
| **Moderate** | Inflammatory infiltrate area found 1-2mm | Epithelial layering in not more than 25% of the epithelium juxtaposed to inflammation in area evaluated | 2 |
| **Severe** | Inflammatory infiltrate area located >2 mm<sup>2</sup> | Epithelial layering in more than 25% of the epithelium juxtaposed to inflammation in area evaluated | 3 |

**Supplemental table 2.**
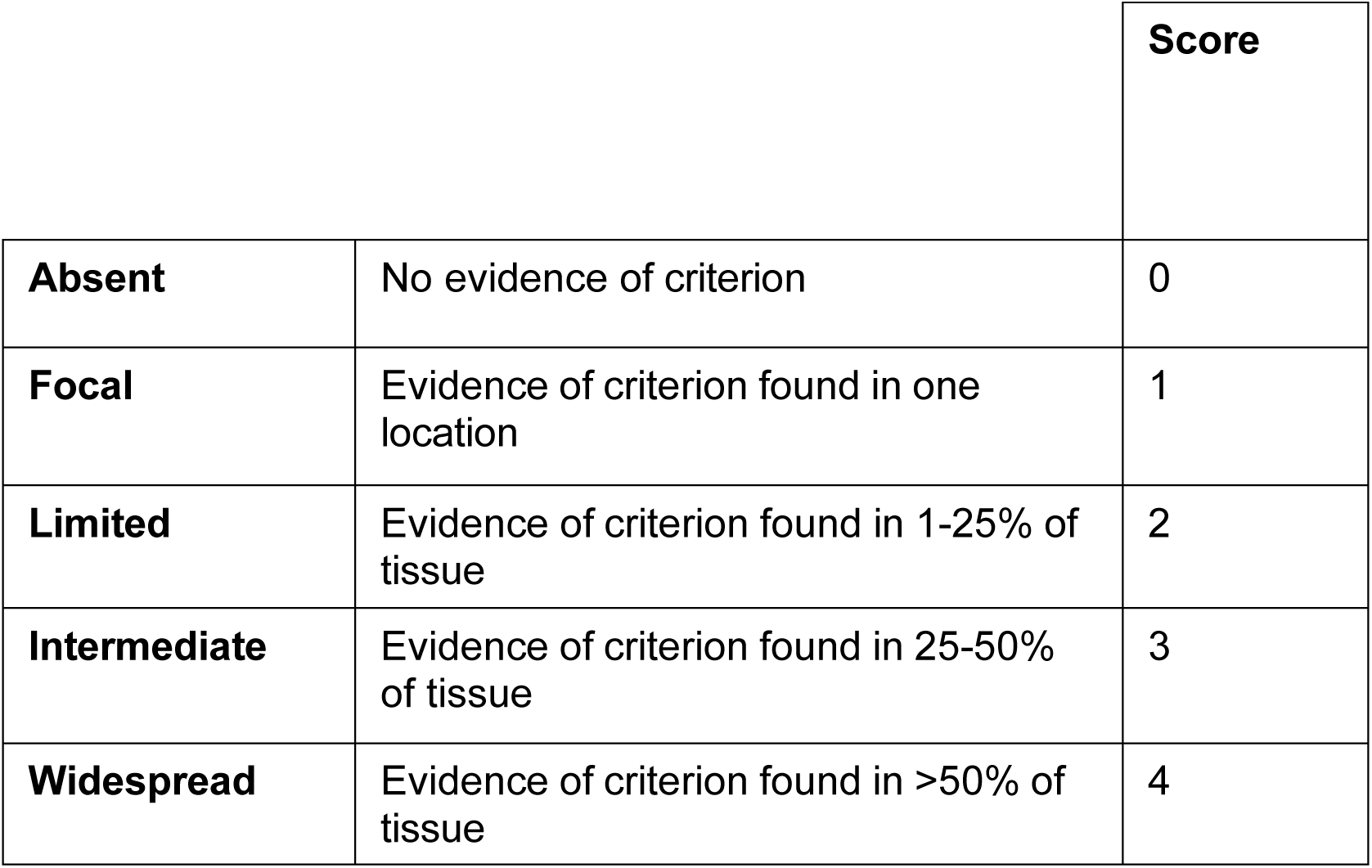
Score system used to quantitate focality of pathology.

|  |  | **Score** |
| --- | --- | --- |
| **Absent** | No evidence of criterion | 0 |
| **Focal** | Evidence of criterion found in one location | 1 |
| **Limited** | Evidence of criterion found in 1-25% of tissue | 2 |
| **Intermediate** | Evidence of criterion found in 25-50% of tissue | 3 |
| **Widespread** | Evidence of criterion found in >50% of tissue | 4 |

**Supplementary Data Sheet X.**
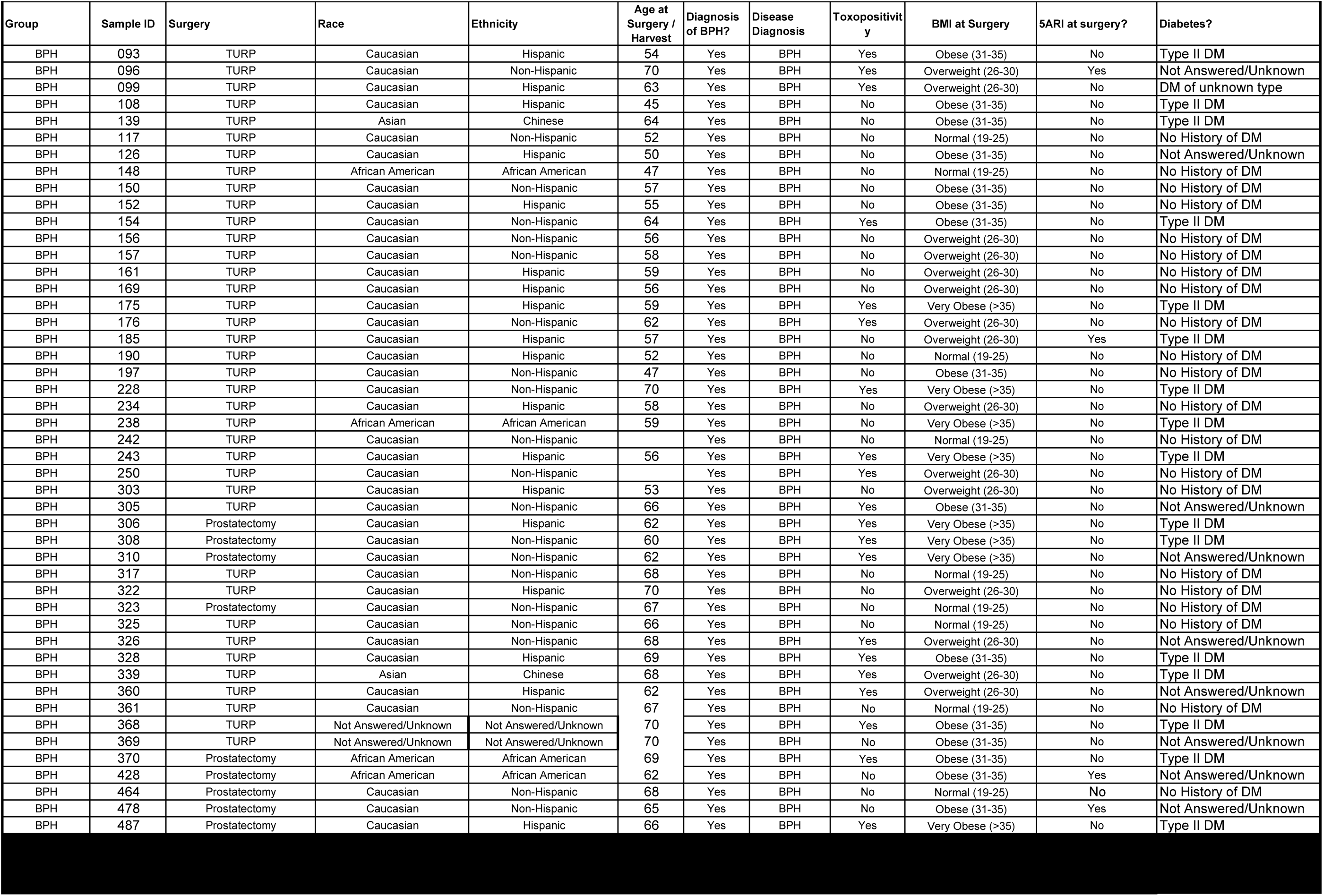
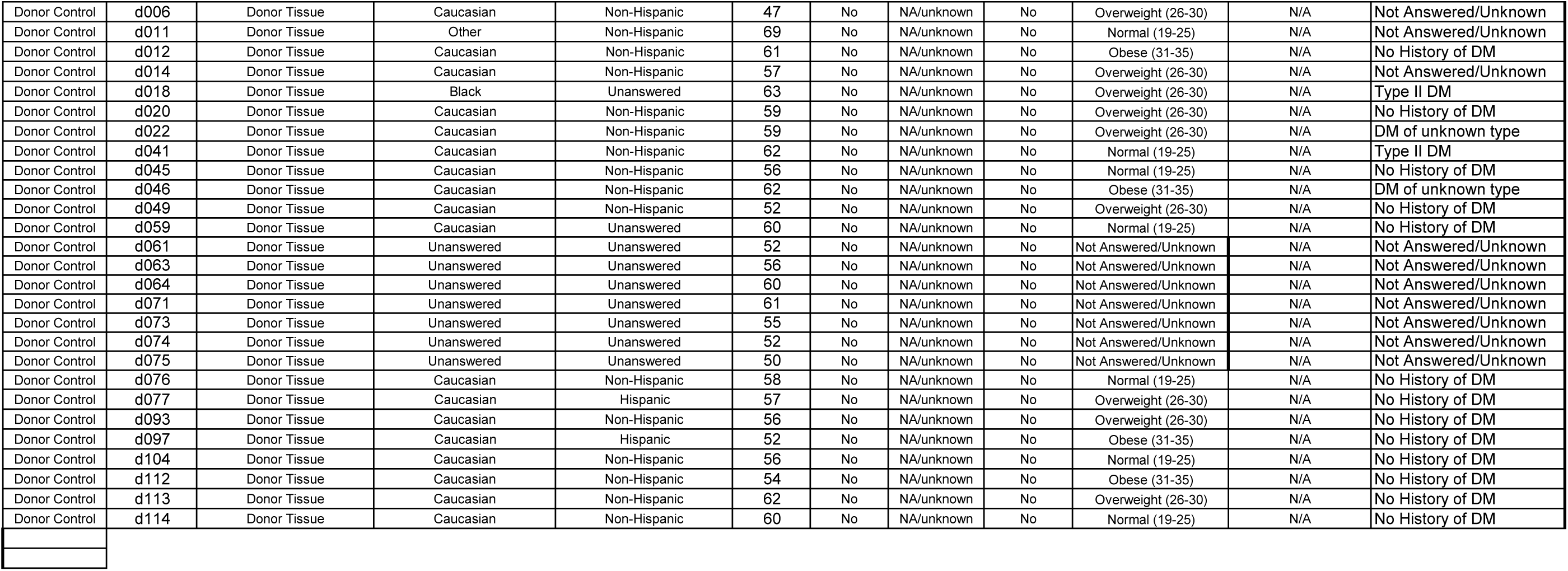
Summary of Human patient characteristics used for Toxoplasma identification and histology studies. General patient and prostate-specific characteristics are indicated for all patients included in used for Toxoplasma serum prevalence and Immunoflorescence staining. Information was gathered via medical record chart review and reflectsclincal information immediately prior to, or at the time of surgery to remove prostatic tissue.

## Notes

**CONFLICTS OF INTEREST:** None.

### Competing Interest Statement

The authors have declared no competing interest.

